# Dimerization and autophosphorylation of the MST family of kinases are controlled by the same set of residues

**DOI:** 10.1101/2023.03.09.531926

**Authors:** Kyler A. Weingartner, Thao Tran, Katherine W. Tripp, Jennifer M. Kavran

**Affiliations:** Department of Biochemistry and Molecular Biology, Bloomberg School of Public Health, Johns Hopkins University, Baltimore, Maryland; Department of Biophysics and Biophysical Chemistry, School of Medicine, Johns Hopkins University, Baltimore, Maryland; The T.C. Jenkins Department of Biophysics, Krieger School of Arts and Sciences, Johns Hopkins University, Baltimore, Maryland; Department of Oncology, School of Medicine, Johns Hopkins University, Baltimore, Maryland

**Keywords:** Hippo pathway, MST kinase, structure-function, protein kinases

## Abstract

The Hippo pathway controls tissue growth and regulates stem cell fate through the activities of core kinase cassette that begins with the Sterile 20-like kinase MST1/2. Activation of MST1/2 relies on *trans*-autophosphorylation but the details of the mechanisms regulating that reaction are not fully elucidated. Proposals include dimerization as a first step and include multiple models for potential kinase-domain dimers. Efforts to verify and link these dimers to *trans*-autophosphorylation were unsuccessful. We explored the link between dimerization and *trans*-autophosphorylation for MST2 and the entire family of MST kinases. We analyzed crystal lattice contacts of structures of MST kinases and identified an ensemble of kinase-domain dimers compatible with *trans*-autophosphorylation. These dimers share a common dimerization interface comprised of the activation loop and αG-helix while the arrangements of the kinase-domains within the dimer varied depending on their activation state. We then verified the dimerization interface and determined its function using MST2. Variants bearing alanine substitutions of the αG-helix prevented dimerization of the MST2 kinase domain both in solution and in cells. These substitutions also blocked autophosphorylation of full-length MST2 and its *Drosophila* homolog Hippo in cells. These variants retain the same secondary structure as wild-type and capacity to phosphorylate a protein substrate, indicating the loss of MST2 activation can be directly attributed to a loss of dimerization rather than loss of either fold or catalytic function. Together this data functionally links dimerization and autophosphorylation for MST2 and suggests this activation mechanism is conserved across both species and the entire MST family.

## INTRODUCTION

The Hippo pathway controls decisions of cell number and cell fate by regulating the activity the transcriptional co-factors YAP/TAZ through a core kinase cassette that includes two kinases MST1/2 and LATS1/2 (1–4). The Hippo pathway is controlled by multiple signals ranging from mechanotransduction to G-protein coupled receptors to cell-cell contacts. A long-standing question was how a diverse set of signals converge on and regulate the activity of the kinase cassette. Activation of the first kinase in the cassette, MST1/2, requires phosphorylation of its activation loop and is largely the result of *trans*-autophosphorylation (5–7). We found cellular events that stimulate MST1/2 activation trigger autophosphorylation by increasing the effective concentration of MST1/2 (8). We then wondered if proximity was sufficient for MST1/2 activation or if additional events were required. Multiple reports suggested a link between kinase-domain dimerization and autophosphorylation for both MST1/2 and its *Drosophila* homolog Hippo (9–11). In fact, MST2 kinase-domains dimerize in solution, but the weak affinity (*K*_*d*_ = 36μM) made the physiological relevance difficult to rationalize. Considering activating events increased the effective concentration of MST1/2 in cells that could, in turn, promote kinase-domain dimerization, we revisited the possible connection between dimerization and autophosphorylation for MST1/2.

Two different dimers have been proposed for MST1/2 and Hippo, but attempts to identify residues required for both functions proved unfruitful and called into question the biological significance of the proposed dimers (9,11). We wondered whether related kinases could inform on the mechanisms of MST1/2 activation. MST1/2 is part of the Sterile 20-like (MST) kinase family that, despite varying biological roles, are each activated by *trans*-autophosphorylation (6,12,13). This family likely shares a conserved activation mechanism owing to the high level of sequence conservation of the kinase-domains (46-88% identity). The family can be further subdivided based on the presence of additional domains (14–16). All start with a kinase domain but the GCK-II sub-family, comprised of MST1 (STK4) and MST2 (STK3), is followed by a linker and a C-terminal SARAH domain while the GCK-III sub-family, comprised of MST3 (STK24), MST4(STK26), and STK25 (SOK1/YSK1), is followed by an alternate C-terminal dimerization domain and bind MO25 proteins(16–18). Studies on the activation of the related kinase MST4, which undergoes autophosphorylation when bound to the activating protein MO25, found the kinase-domain to also weakly associate in solution with a *K*_*d*_ in the tens of micromolar and also arrived at two different models for dimerization, and attributed the differences in models to representing different steps of autophosphorylation (13,19).

We set out to understand whether or how dimerization contributes to autophosphorylation MST2 and the entire MST family. We started by analyzing crystal lattice contacts of MST kinase-domains to identify potential dimers compatible with *trans*-autophosphorylation. Our analysis identified eight potential dimers, each of which was mediated by the activation loop and the αG-helix on the C-lobe. These dimers clustered into three groups based on both the relative arrangement of the kinase-domains within the dimers and the activation state of the kinase or presence of binding partners. We then validated these structural observations with functional studies of the αG-helix using MST2. Variants with alanine substitutions of the αG-helix did not dimerize either *in vitro* or in cells, establishing the αG-helix is required for dimerization. The same substitutions impaired autophosphorylation of both full-length protein in cells and of kinase-domains in a purified system. Equivalent substitutions in Hippo also impaired activation in cells. The lack of autophosphorylation, which requires dimerization of the kinase-domains, can be attributed to substitution of these residues, that neither disrupted folding nor substrate phosphorylation. Together our data links dimerization to autophosphorylation of MST2 through the αG-helix and suggests this activation mechanism is conserved across the MST family and across species.

## RESULTS

### A conserved region mediates dimerization of MST kinase-domains

Given the sequence conservation between the kinase-domains of the MST family, we and others reasoned these kinases likely share a conserved mechanism of activation(13) (***Figure 1***). We set out to understand the nature of kinase-domain dimerization for the MST family and perhaps identify conserved features that may guide activation of the family and thus provide insight into the specifics of MST2 activation. To understand how these kinase-domains dimerize and possibly resolve differences between proposed models, we identified all unique observations of kinase-domain pairs generated by crystal symmetry of previously determined structures of the MST family (MST1-4 and STK25) (11,13,19–28). This analysis identified 73 unique kinase-domain pairs (***Table 1***).

**Table 1.** Structures used in kinase-domain dimer analysis * kinase-domain dimers are referred to by the nomenclature “dimer number_protein_PDBID”

| Dimer Numbering | Protein | Activation loop phosphorylation | PDBID | Reference |
| --- | --- | --- | --- | --- |
| 1 – 5 | MST1 | + | 3COM | (20) |
| 6 – 9 | MST1 | + | 6YAT | (21) |
| 10 – 27 | MST2 | - | 4LG4 | (11) |
| 28 – 33 | MST2 | + | 5DH3 | (22) |
| 34 – 40 | MST2 | - | 4LGD | (11) |
| 41-42 | MST2 | - | 6AO5 | (23) |
| 43 – 46 | MST3 | + | 3A7J | (24) |
| 47 | MST3 | - | 3ZHP | (25) |
| 48 – 51 | MST3 | + | 4QMM | (26) |
| 52 – 57 | MST3 | + | 3A7G | (24) |
| 58 | MST3 | - | 4O27 | (27) |
| 59 – 66 | MST4 | - | 3GGF | (13) |
| 67 – 69 | MST4 | - | 4FZA | (19) |
| 70 – 72 | STK25 | + | 2XIK | (28) |
| 73 | STK25 | - | 4NZW | (27) |

**Figure 1.**
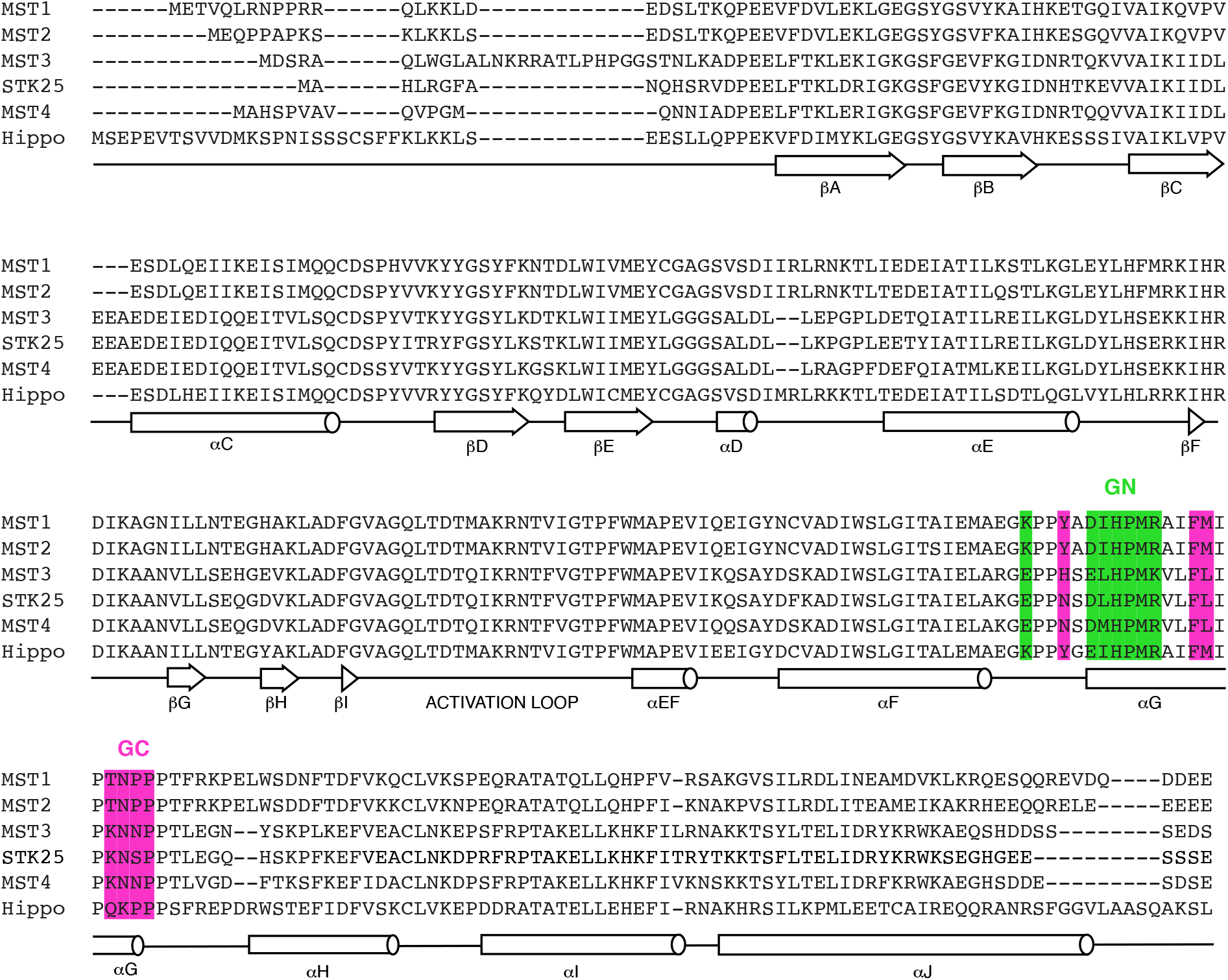
MST sequence alignment. Sequence alignment of Sterile-20 like kinase-domains. A secondary structure assignment is shown below the alignment and is based on a validated multiple-sequence alignment of kinase-domains(44). Substitutions corresponding to GN and GC are colored green and pink, respectively.

We first asked if the same kinase-domain dimer mediated lattice contacts in multiple crystal forms since biologically relevant interfaces are more likely to mediate crystal contacts than non-physiological ones (29,30). A well-known example of the same biological interface mediating contacts in different lattices is the asymmetric kinase-domain dimer of Epidermal Growth Factor Receptor (EGFR) (31). We compared the structure of each kinase-domain pair to every other pair by iterative superpositions and found no conserved arrangement among all the pairs or any subgroup (***Table S1, S2***). We then asked if any kinase-domain pairs were compatible with *trans*-autophosphorylation. Rather than identifying dimers correlated with a specific step in *trans*-autophosphorylation, we used two criteria that could describe any step in the reaction. First, potential dimers must bury sufficient surface area to be considered a biologically relevant interface. In crystal lattices, biological assemblies are larger than non-physiological ones and typically bury at least 500Å_2_ of surface area (29,30). We calculated the surface area buried in each kinase-domain pair (***Table S3***) and found nearly half satisfied this requirement. Second, within a potential dimer the kinase-domains should be poised for *trans*-autophosphorylation, specifically the activation loop of one kinase must be able to extend into the catalytic cleft of its partner. We calculated whether each activation loop could theoretically extend into the catalytic cleft of its partner and found eleven were properly poised pairs (***Table S3***). From our initial set, eight pairs satisfied both requirements (1_3COM_MST1, 6_MST1_6YAT, 11_4LG4_MST2, 12_4LG4_MST2, 30_5DH3_MST2, 34_5DH3_MST2, 59_3GGF_MST4, and 69_4FZA_MST4) (***Table 2, Figure 2, S1***). These dimers represented the spectrum of structures analyzed, containing representative from three out of the five MST family members and both phosphorylated and unphosphorylated kinases. This set included the four previously proposed dimers as well as four novel dimers (9,11,13,19).

**Figure 2.**
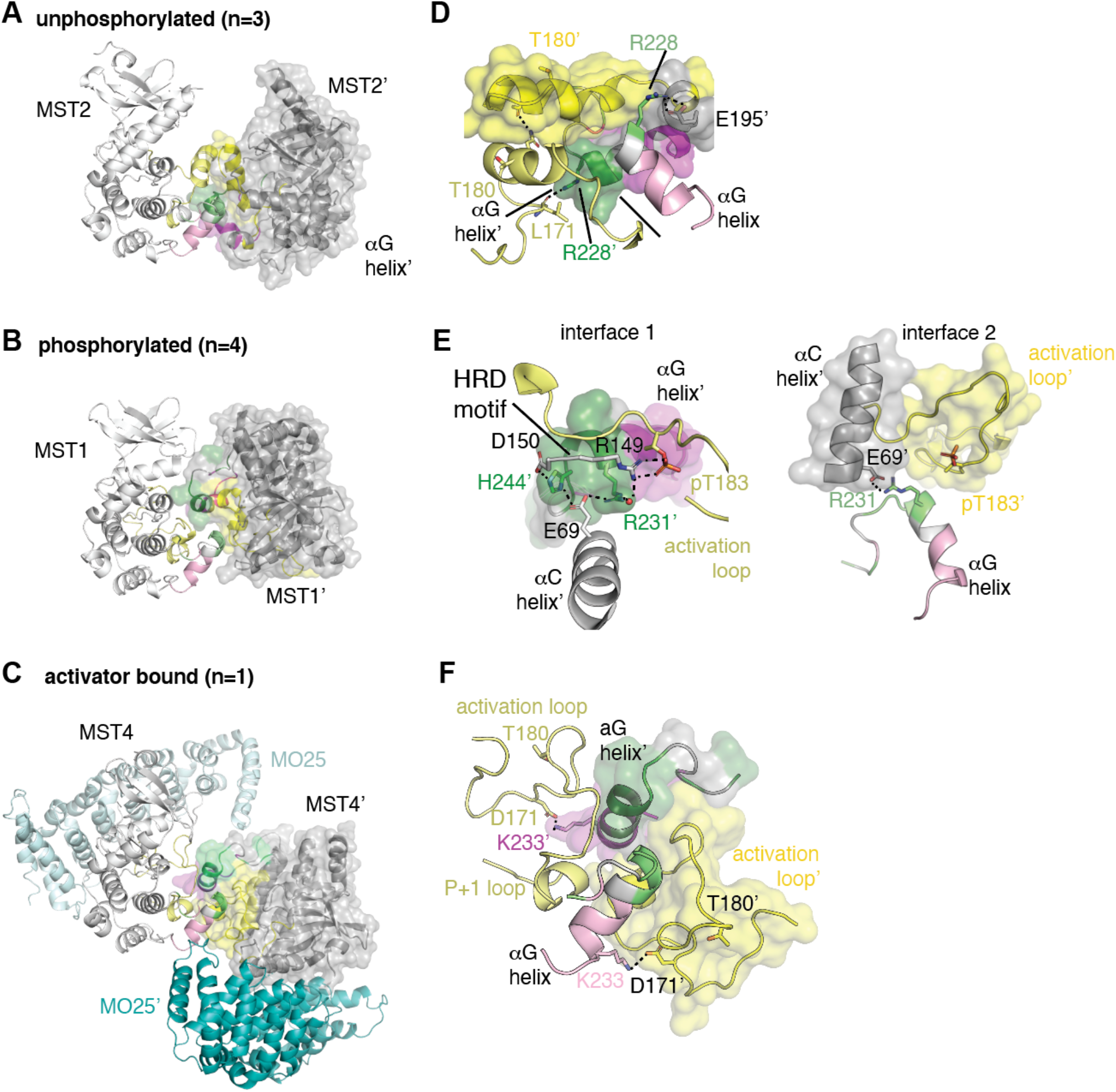
Three conformations of MST kinase-domain dimers. Gallery of the highest resolution structure from each group including (A) an unphosphorylated dimer (11_4LG4_MST2), (B) a phosphorylated dimer (1_3COM_MST1), and (C) an unphosphorylated dimer bound to the activator MO25 (teal) (69_4FZA_MST4). (D,E,F) A close up of each dimer interface. For each panel, one kinase is shown in white cartoon while the other in both gray cartoon and surface. The N-terminal and C-terminal regions of the αG-helix are colored in shades of pink or green, respectively, and activation loops in shades of yellow. Residues mediating inter-kinase interactions are displayed in sticks.

**Table 2.** Kinase-domain dimers that satisfy the requirements for trans-autophosphorylation

| Dimer | Buried Surface Area (Å<sup>2</sup>) | Properly Poised | αG-mediated |
| --- | --- | --- | --- |
| 1_3COM_MST1<sup>c</sup> | 798 | Yes | Yes |
| 6_6YAT_MST1 | 750 | Yes | Yes |
| 11_4LG4_MST2<sup>c</sup> | 831 | Yes | Yes |
| 12_4LG4_MST2 | 634 | Yes | Yes |
| 30_5DH3_MST2 | 904 | Yes | Yes |
| 31_5DH3_MST2 | 746 | Yes | Yes |
| 59_3GGF_MST4<sup>c</sup> | 1001 | Yes | Yes |
| 69_4FZA_MST4<sup>c</sup> | 1062 | Yes | Yes |

<sup>a</sup> “Yes” indicates at least one chain satisfies requirements
<sup>b</sup> “Yes” equates to at least one residue in the indicated region is buried at the dimer interface.
<sup>c</sup> Indicates a previously identified kinase-domain dimer

The dimer interface of each of these dimers was mediated by the same regions, the activation loop and αG-helix (***Figure 2, S1***). The relative orientation of the kinases within these dimer, however, varied clustered into three groups based on the relative position of the kinase-domains as well as the phosphorylation state of the kinase-domain or presence of activating proteins (***Figure 2, S1***). One group contains the four phosphorylated dimers (1_3COM_MST1, 6_6YAT_MST1, 30_5DH3_MST2, and 31_5DH3_MST2), another the three unphosphorylated dimers (11_4LG4_MST2, 12_4LG4_MST2, and 59_3GGF_MST4), and the final a single dimer bound to the activator protein MO25 (69_4FZA_MST4). Dimers of MST2 and MST4 appear in both the phosphorylated and unphosphorylated clusters, perhaps revealing the range of structural space these kinase sample during *trans*-autophosphorylation (***Figure S1***).

We analyzed the interface of the highest resolution member of each group to understand the specific interactions mediating dimerization of each group (***Figure 2)***. For the unphosphorylated kinase domain dimer, the αG-helices form a single docking site that engages both activation loops (***Figure 2A***). In this arrangement, the site of phosphorylation (T180, MST2 numbering) is facing away from the interface suggesting it may represent a pre-catalytic state. The phosphorylated kinase-domain dimer, which is the only asymmetric dimer, has two interfaces that each engage several structural hallmarks of the active kinase-domain conformation (***Figure 2B***). In one interface, the αG-helix of one kinase binds the activation loop (pT183, MST1 numbering), HRD motif (R149, D150), and αC-helix (E69) of its partner through a series of non-bonded contacts and an extended hydrogen bonding network. In the other interface, the other αG-helix engages the αC-helix of its partner kinase forming a surface on which the activation loop of its partner kinase docks. In both interfaces E69 of the αC-helix forms a salt bridge with R231 of the αG-helix. For the MST4 kinase dimer bound to MO25, the dimerization is mediated by the αG-helices as well as the activation and P+1 loops and a two equivalent inter-kinase salt bridges using K233 (MST4 numbering) and D171. The activator protein does not make any contribution to the dimer interface (***Figure 2C***).

### The αG-helix mediates kinase domain dimerization

We wanted to validate the interfaces in observed in the structures and determine the function of the αG-helix in dimerization and activation of MST2. To avoid disrupting the fold of the kinase-domain, we made two variants in which we substituted a set of surface residues that comprised either the N-terminal (MST2-K_GN_) or C-terminal (MST2-K_GC_) region of the αG-helix with alanine. We did not make any substitutions in the activation loop as they would likely impair the function of the kinase-domain. The variants were expressed and purified similar to wild-type. To ensure the variants had the same fold as wild-type we compared the secondary structures of MST2-K_WT_, MST2-K_GN_, and MST2-K_GC_ using far-UV circular dichroism (CD) (***Figure 3***). Each of the three far-UV CD spectra have a well-defined minima at 208nm and a shallower minima at 222nm indicating a mix of α-helical and -sheet that is consistent with the known secondary structure composition of the kinase-domain. Within error, the spectra of the alanine variants overlay with wild-type indicating that the substitutions did not disrupt the kinase-domain fold.

**Figure 3.**
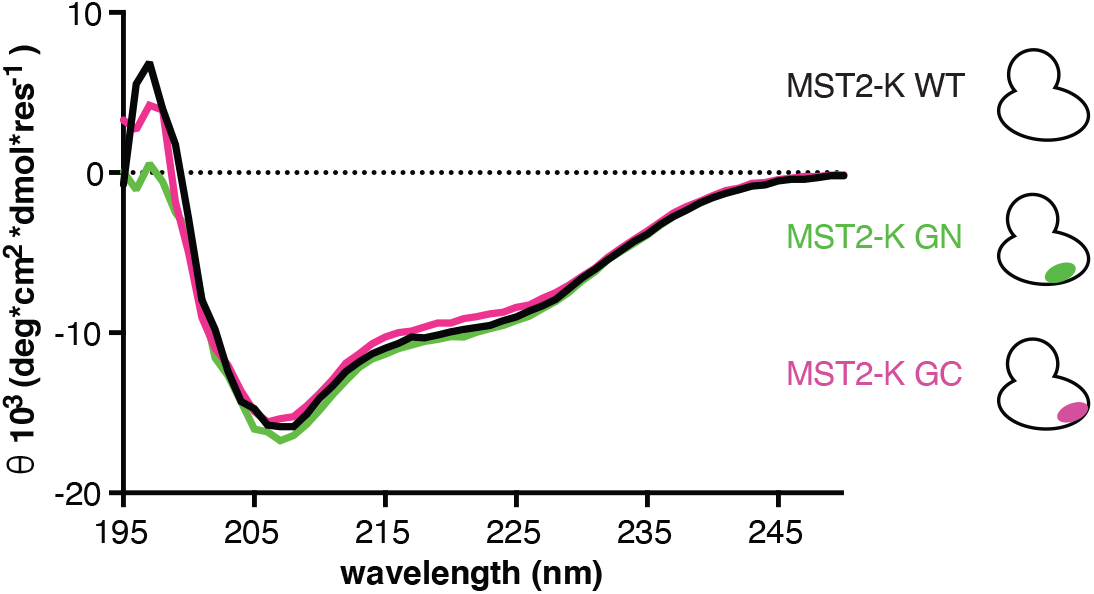
Substitutions of αG-helix do not disrupt secondary structure. Overlay of far-UV CD spectra for MST2-K_WT_ (black), MST2-K_GN_ (green) and MST2-K_GC_ (pink).

Then we directly monitored dimerization of the MST2 kinase-domains using sedimentation analytical ultracentrifugation (AUC) and compared the solution behavior of wild-type (MST2-K_WT_) to both αG-helix variants, (MST2-K_GN_ and MST2-K_GC_). We computed the apparent sedimentation coefficient distributions, g(s*), for all proteins over a range of concentrations. For MST2-K_WT_, upon increasing concentration we observed a shift from lower to higher apparent Svedberg coefficient indicative of monomer-dimer equilibrium. For MST-K_GN_ and MST-K_GC_, no shift was observed over the protein concentration analyzed (***Figure 4A***) suggesting these variants, which have a disrupted

**Figure 4.**
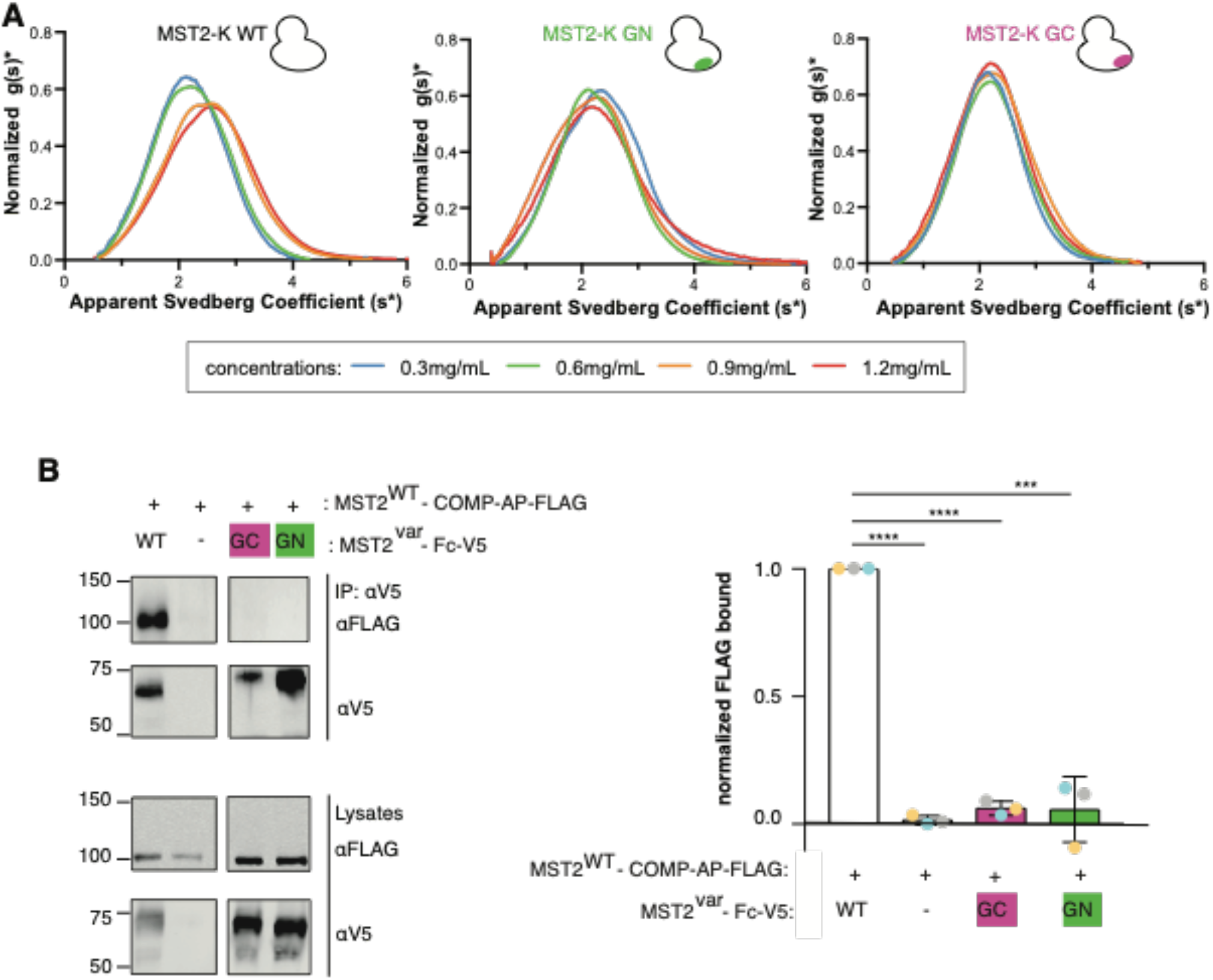
Substitution of the αG-helix prevents dimerization of MST2 kinase-domains. (**A**) Sedimentation coefficient distributions of MST2-K_WT_,MST2-K_GN_, and MST2-K_GC_ displayed as g(s*) plots. Colors of the distribution curves correspond to the concentration of the proteins, as indicated. (**B**) HEK293 cells transiently transfected as indicated, and complexes isolated from lysates following avidity-based immunoprecipitation with V5 antibody. Protein expression and complex formation were monitored by Western blot. (*Left*) A representative set of blots are shown, panels are from same blot but cropped for clarity. Full westerns and replicates are in ***Figure S2***. (*Right*) Band intensities of bound FLAG-tagged proteins were quantified ((bound-background)/normalization). The bar graph represents the mean from three experiments, and the errors bars the standard deviation. Significant differences were calculated using an unpaired *t*-test (***, *p* ≤ 0.0008, ****, *p* ≤ 0.0001).

### αG-helix, were monomeric

We then asked if these substitutions disrupted MST2 kinase-domain dimerization in cells. We first attempted to detect oligomerization between MST2 kinase-domains using standard immunoprecipitation assays. While we could detect a complex between differentially tagged full-length MST2, which form a high affinity complex owing to dimerization of the C-terminal SARAH-domains (32,33), no interactions were detected between differentially tagged kinase-domains (***Figure S2***) which is consistent with the reported weak disassociation constant.

Avidity-based approach are able to detect weaker interactions that straight immunoprecipitations owing to the use of oligomeric tags, so we employed that strategy to detect interactions between MST2 kinase-domains (34). For the bait, we used a MST2-K fused to the dimeric Fc protein with a V5 tag for detection. For the prey, MST2-K was fused to the pentameric cartilage outer matrix protein (COMP) and a FLAG tagged added for detection and alkaline phosphatase (AP) for a mass tag to help distinguish the different variants on Western blots. With this approach we successfully detected binding of avidity-tagged wild-type MST2-K (***Figure 4B, S3***). No interactions, however, were detected for either variants bearing substitutions of the αG-helix (MST2-K_GN_ and MST2-K_GC_) (***Figure 4B***) demonstrating that disruption of a single αG-helix is sufficient to prevent oligomerization. Together these data suggest the αG-helix mediates dimerization of MST2 kinase-domains.

### Dimerization is required for autophosphorylation

We wondered if the residues required for dimerization were also required for autophosphorylation. Phosphorylation of the activation loop of MST2 was monitored by Western blot in HEK293 cells transiently transfected with plasmids encoding either wild-type MST2-FL (MST2-FL_WT_), MST2-FL variants bearing substitutions of αG-helix (MST2-FL_GN_ or MST2-FL_GC_), or the kinase-inactivating substitution (MST2-FL_D146N_) (***Figure 5A***). No activation loop phosphorylation was detected for either MST2-FL_GN_, MST2-FL_GC_, or MST2-FL_D146N_ but was detected for MST2-FL_WT_. Disruption of the αG-helix impairs autophosphorylation of MST2.

**Figure 5.**
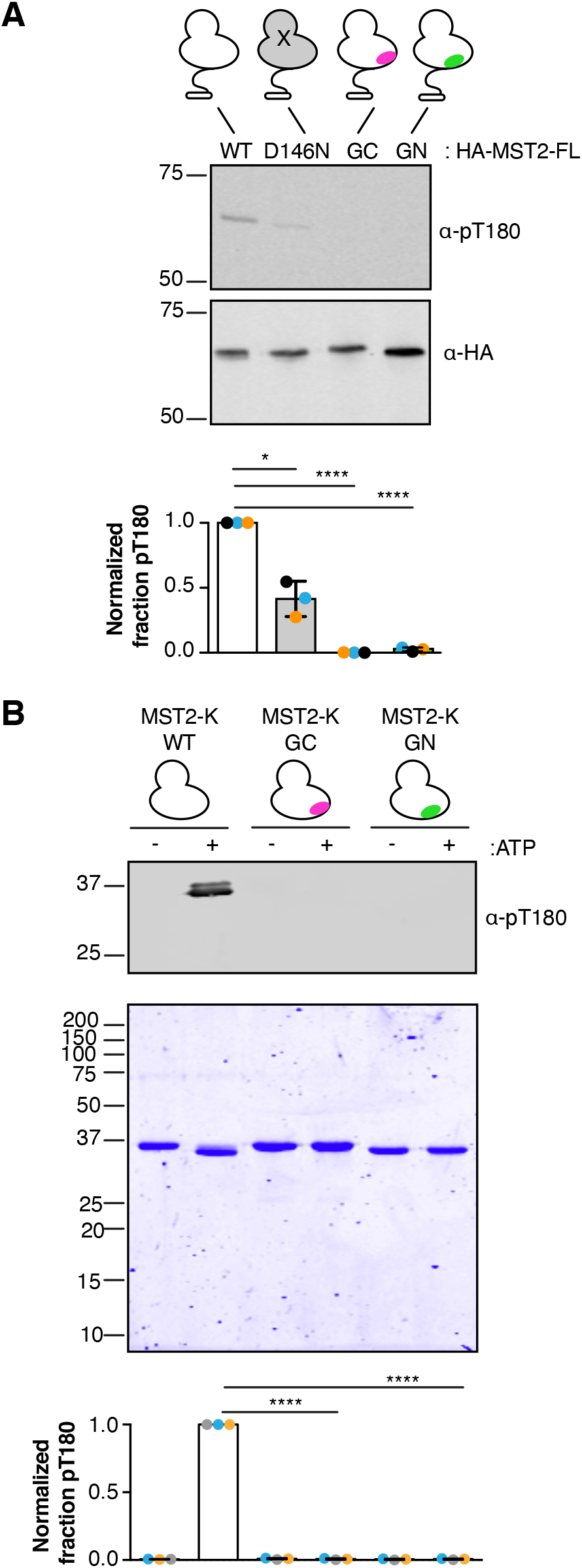
Dimerization deficient MST2 variants do not undergo autophosphorylation. (A) Cell lysates from HEK293 cells transiently transfected with plasmids encoding variants of MST2-FL, as indicated, were analyzed by Western blot. The experiment was performed three times, and a representative set of blots are shown. Band intensities from each replicate were quantified, and the fraction of phosphorylated MST2 determined. The normalized values are plotted, with colors corresponding to sets of replicates, below each lane. Bar graphs represent the average value and error bars corresponding to the standard deviation. (B) Purified, unphosphorylated MST2-K variants were incubated in the presence or absence of ATP. Autophosphorylation was monitored by Western blot, and total protein tracked by Coomassie stained SDS-PAGE gel. The experiment was performed three times, and a representative result is shown. Plotted below the corresponding lanes of the gels are each individual data point, colored coded by replicate. The mean values are shown as a bar graph, and the error bars the standard deviation. Significant differences were calculated using an unpaired *t*-test (*, *p* ≤ 0.002; ****, *p* ≤ 0.0001).

To ensure the lack of autophosphorylation for MST2-FL_GN_ and MST2-FL_GC_ could be directly attributed to changes in the kinase-domain rather than interactions with other domains or proteins found in a cell, we analyzed autophosphorylation of MST2-K_WT_, MST2-K_GN_ and MST2-K_GC_ *in vitro*. mWe expressed and purified three unphosphorylated variants corresponding to wild-type (MST2-K_WT_) and variants with a substituted αG-helix (MST2-K_GN_ or MST2-K_GC_), incubated each with ATP and Mg_+2_, and monitored phosphorylation by Western blot (***Figure 5B***). MST2-K_WT_ autophosphorylated, but MST2-K_GN_ or MST2-K_GC_ did not suggesting the loss of autophosphorylation is a direct consequence of disruption of the αG-helix.

### MST2 variants are catalytically active

We also wanted to determine if the kinase-domain variants retained catalytic function so the observed lack of autophosphorylation could be attributed to disruption of dimerization rather than a simple loss of catalytic activity. We monitored the ability of wild-type and variant kinase-domains to phosphorylate a substrate. We made activated MST2-K variants by incubating unphosphorylated MST2-K_WT_, MST2-K_GN_, MST2-K_GC_ or a catalytically inactive variant, MST2-K_D146N_, with ATP and phosphorylated MST2-FL, after which MST2-FL was removed by affinity purification. The phosphorylated MST2-K variants were incubated with a known substrate, MOB1A, and phosphorylation of MOB1A monitored by Western blot (***Figure 6***). MST2-K_D146N_ serves as a negative control to determine if any active MST2-FL carried through to the substrate phosphorylation assay. Minimal phosphorylation of MOB1A was observed for MST2-KD146N, thus ensuring any phosphorylation of MOB1A can be attributed to the MST2 variant tested. MST2-K_WT_, MST2-K_GN_, and MST2-K_GC_ each phosphorylated MOB1A demonstrating that substitution of the αG-helix did not destroy catalytic function. We note the different levels of MOB1A phosphorylation likely reflect the varying levels of active MST2 in each reaction, as judged by the fraction of MST2 phosphorylated on pT180 (***Figure 6***), which would have different rates of substrate phosphorylation.

**Figure 6.**
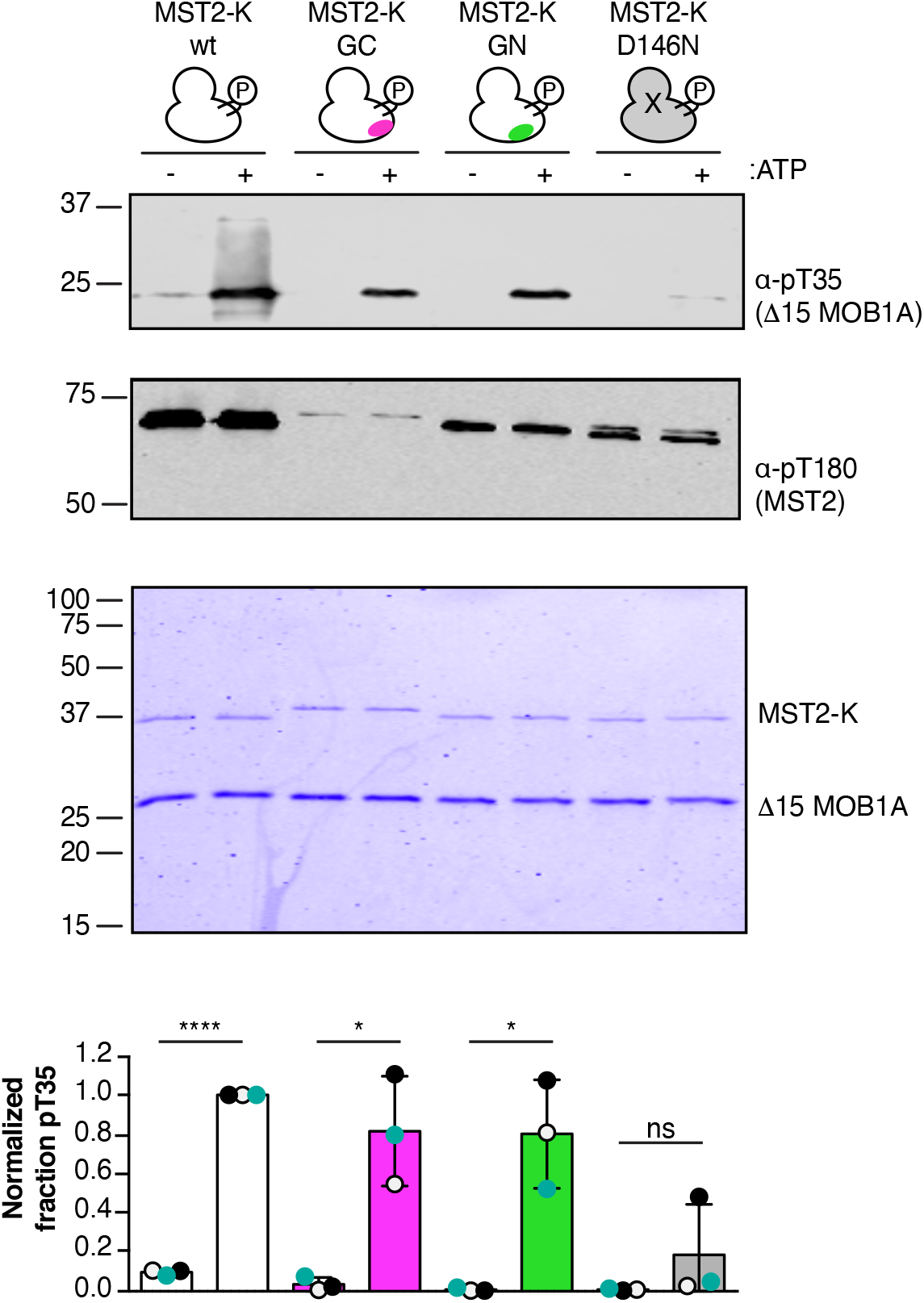
Substitution of αG-helix does not disrupt substrate phosphorylation. (*Top*) Indicated variants of phosphorylated MST2-K were incubated with ι115 MOB1A in the presence or absence of ATP. Phosphorylation of ι115 MOB1A was monitored by Western blot using a phospho-specific antibody (α-pT35) and by Coomassie stained SDS-PAGE. Levels of autophosphorylation of MST2-K variants was monitored by α-pT180. The experiment was performed three times, and a representative set of gels are shown. (*Bottom*) Band intensities were quantified and the normalized fraction of phosphorylated Mob1 calculated. Individual data points are shown as circles and color coded according to replicate. The bar graph represents the mean from three replicates, and error bars the standard deviation. Significant differences were calculated using a paired *t*-test (ns, no significant difference; *, *p* ≤ 0.05; ****, *p* ≤ 0.0001)

### Equivalent residues mediate autophosphorylation of Hippo

We analyzed the conservation of the αG-helix in Hippo and found it was absolutely conserved barring one residue (T235 in MST2 is Q250 in Hippo) (***Figure 1***). Given this conservation, we wondered if autophosphorylation of Hippo relied on equivalent residues as MST2. HEK293 cells were transiently transfected with plasmids encoding either wild-type Hippo (Hippo-FL_WT_), catalytically inactive Hippo (Hippo-FL_K51R_), or variants bearing substitutions equivalent to either the N-terminal or C-terminal half of the αG-helix surface (Hippo-FL_GN_, Hippo-FL_GC_), and autophosphorylation monitored by Western blot (**Figure 7**). We observed robust phosphorylation for Hippo-FL_WT_ but not Hippo-FL_K51R_ demonstrating the phosphorylation observed is from autophosphorylation. Substitution of the αG-helix in Hippo prevented autophosphorylation suggesting this function of this helix is conserved across species.

**Figure 7.**
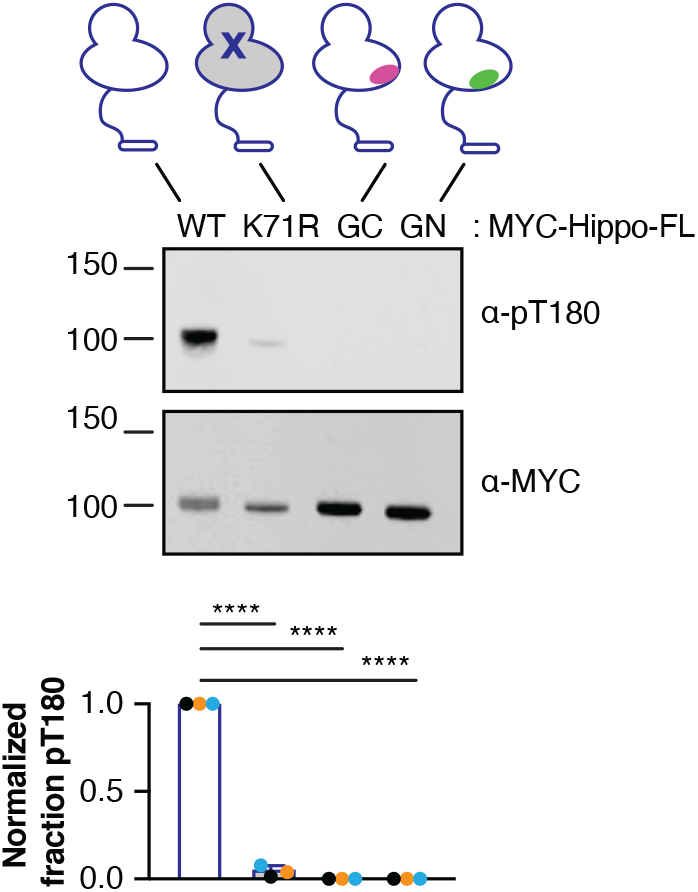
Disruption of αG prevents autophosphorylation of Hippo. A representative set of western blots detecting protein expression (α-MYC) or activation loop phosphorylation (α-pT180) of MYC-Hippo-FL variants following transient transfection of HEK293 cells. The experiment was performed three times, and the band intensities from each replicate were quantified. Plotted below each lane are the data from each replicate displayed as the normalized fraction of phosphorylated protein. Bar graphs represent the average value and error bars corresponding to the standard deviation. Significant differences were calculated using an unpaired *t*-test (****, *p* ≤ 0.0001).

## DISCUSSION

We set out to understand the role of kinase-domain dimerization in activation via *trans*-autophosphorylation for the MST family and potentially resolve the differences between the dimer proposed dimerization models. Our data provides clear evidence that a conserved set of residues mediate dimerization associated with *trans*-autophosphorylation for the entire MST family and, by using MST2 as a model system, validate the structural models of dimerization and provide a functional link between dimerization and autophosphorylation. Our work rationalizes previous results and further suggests this mode of dimerization may be evolutionarily conserved for MST kinases.

Establishing a role for the αG-helix in both dimerization and autophosphorylation of MST2 furthers our understanding molecular mechanism of MST2 activation. Earlier work suggested activating events that increased the proximity of MST2 triggered autophosphorylation (8). This model can now be modified to increased proximity promotes autophosphorylation by promoting dimerization that is required for autophosphorylation. This model predicts regulatory proteins that bind to the residues required for dimerization would block activation of MST kinases. Our data also demonstrate that requirements for autophosphorylation are different than substrate phosphorylation as MST2 variants unable to undergo autophosphorylation could still phosphorylate a protein substrate (**Figure 5**,**6**).

Our structural analysis revealed the MST family shares a conserved mode of dimerization (***Figure 2, Figure S1***). The interface of each potential *trans*-autophosphorylation dimer was mediated by the αG-helix, a highly conserved region between human GCKII and GCKIII families (**Figure 1**) (50% identity). We validated the role of αG-helix when we found substitution of these residues disrupted both dimerization and autophosphorylation of MST2 (**Figures 4**,**5**); the role of these residues in activation had been previously validated for MST4 (19). We also find that these same residues are required for autophosphorylation of Hippo, suggesting this activation mechanism is conserved across species (***Figure 7***). The identity of this region is not maintained in other GCK kinase families (0% identity) suggesting dimerization modes vary between kinase families.

We did not anticipate that multiple dimers would be compatible with *trans*-autophosphorylation. While initially perplexed by the lack of a common dimer, several lines of reasoning supported activation being compatible with multiple conformations. A fundamental difference between MST and ErbB kinases is the nature of kinase domain activation. ErbBs do not require autophosphorylation for activation so the kinase dimer must be maintained for activity (31). In contrast, activity of MST family requires activation loop phosphorylation, but once phosphorylated the *trans*-autophosphorylation dimer is no longer required and must dissemble to accommodate substrate binding. This *trans*-autophosphorylation dimer represents an enzyme:substrate complex that by nature is a less stable, has a shorter half-life than typical protein:protein interactions, and must disassociate to allow phosphorylation of other substrates by the now activated kinase. Further, the weak affinity of the kinase-domain dimer of the MST2 kinase domain equates to a significantly lower energetic requirement for complex assembly than would be required by stronger affinity complexes. This lower energetic barrier could be satisfied by multiple conformations.

We identified 8 ensemble dimers that clustered into three groups based the relative arrangement of the kinase-domains within the dimers and aligned with the presence or absence of activation loop phosphorylation or activator proteins (***Figure 2, S1***). It is, indeed, tempting to speculate these groups represent pre- and post-catalytic states; all the kinases-domains in the phosphorylated group adopt the active conformation and the dimer interface engages multiple regulatory elements (αC-helix, HRD motif, and the activating site of phosphorylation on the activation loop) that stabilize that active conformation. In contrast, the kinases in the unphosphorylated group adopt a variety of partially active conformations. The structural variation within this ensemble may represent the structural space sampled by a kinase during *trans*-autophosphorylation. In the future, experiments looking at the structural dynamics of *trans*-autophosphorylation should investigate the repacking of the dimer interface and the rearrangements within the kinase that occur during *trans*-autophosphorylation. Our work suggests a conserved interface mediates dimerization and activation for the entire MST family and it will be interesting going forward investigating how molecular mechanisms of activation vary between kinase families.

### EXPERIMENTAL PROCEDURES

#### Generation of lattice dimers

The Protein Data Bank(35) on 7/5/20 data contained 47 crystal structures of GCKII and GCKIII family members. This list was then pruned to contain only unique crystal forms. By this process we identified 15 unique crystals forms that included representatives from each of the five family members and both phosphorylated or unphosphorylated kinases (***Table 1***). If binding partners or other domains were present in the structures, those components were removed for generation of lattice-mediated kinase-domain pairs. We then manually identified all unique kinase-domain pairs mediated by contacts either within the asymmetric unit or by crystal symmetry using UCSF Chimera(36) resulting in 73 independently observed kinase-domain pairs.

#### Structural Analysis

Both buried surface area calculations and identification of residues buried at the interface were performed in AREAIMOL in CCP4 (37). The criteria for the αG-helix to be mediating dimerization was burial of a single residue. To determine the similarity of kinase-domain dimers, the C-lobe of every kinase was superposed onto each kinase-domain in every other kinase-domain dimer using LSQKAB in CCP4 (37). If the superposition generated an RMSD less than 3Å over the 78 Cα-atoms, then dimers were considered similar. To determine if the kinases were properly positioned, we first modeled the position of ATP in the catalytic cleft of one kinase by superimposing the structure of MST3 bound to AMP-PNP onto each kinase in the pair (PDBID 4QML)(26) and then measured the distance between the γ phosphate and either the start (residue 162, MST2 numbering) or end of the activation loop (residue 185) of the partner kinase. Kinases were deemed properly poised if, assuming the activation loop was fully extended, T180 could be positioned in the activation cleft of its partner.

#### Avidity pull-downs

HEK293T cells were seeded at 0.4*10^6^cells/well in 6-well plates, 24 hours later they were transfected with indicated plasmids, and lysed twenty-four hours afterwards in ice-cold Immunoprecipitation Buffer supplemented with protease and phosphatase inhibitors (20mM Tris pH 8, 150mM NaCl, 1% Nonidet P-40, 10% Glycerol) supplemented with 1mM Phenylmethylsulfonyl fluoride (PMSF), 1mM Na_3_VO_4_, 10mM NaF, 2.5mM Na_2_P_2_O_7_, 1mM β-glycerophosphate, Protease Inhibitor Cocktail (Sigma, #P8849), and Universal Nuclease (Pierce, #88701). Lysates were normalized using the BCA Assay (ThermoFisher, #23225) and then incubated with Protein G resin (homemade) and V5 antibody (Cell Signaling Technologies, Lot 6) for 3 hours at 4_o_C. Resin was collected, washed three times each with IP buffer, and boiled in SDS-loading buffer. For Western blots of the proof-of-concept pulldowns, samples were analyzed by Western blot using primary antibodies that recognized either the V5 (Cell Signaling, Lot 6, diluted 1:1000) or FLAG (Sigma-Aldrich, Lot SLCD3990, diluted 1:1000) epitopes followed by either IRDye 800CW Goat anti-Rabbit (LI-COR, Lot C90220-06, diluted 1:10,000) or IRDye 800CW Goat anti-Mouse (LI-COR, Lot C81106-01, diluted 1:10,000) secondary antibody, respectively. Blots were scanned on an Odyssey Infrared Imaging System (LI-COR). For Western blots of the alanine-substitution variants the same protocol was followed except for the use of an anti-Rabbit HRP-linked secondary antibody (Cell Signaling, Lot 28, diluted 1:10,000) for the V5 antibody. Then blots were incubated with Clarity Max_TM_ Western ECL Substrate (Bio-Rad), and scanned with a GBox Chemi XX9 System (SYNGENE). Band intensity were quantified by ImageJ (38). The amount of normalized, bound protein was calculated by first subtracting the amount of background binding for each variant and then expressing the amount of bound protein as a fraction relative to the amount of bound, wild-type MST2-FL or wild-type MST2-K fused to the COMP avidity tag.

#### Protein expression and purification

DNA encoding residues 1-314 of human MST2 kinase-domain (UniprotKB Q13188) were cloned downstream of a hexa-histidine and SUMO tags into either a modified pBAD4 plasmid(39) for MST2-K_wt_ and MST2-K_D146N_, or pRSF-Duet (EMD-Millipore, MA) for MST2-K_GN_ and MST2-K_GC_. MST2-K_GN_ had residues (Y221, F231, M232, T235, N236, P237, P238) substituted to alanine and MST2-K_GC_ had these residues (K218, D223, I224, H225, P226, M227, R228) substituted with alanine. For expression, MST2 variants were transformed into T7 express cells (New England BioLabs, MA) along with a plasmid encoding maltose binding protein (MBP) λ-Phosphatase. Bacterial cultures were grown at 37°C until OD600 of 2.0, and protein expression induced with 0.25mM Isopropyl β-d-1-thiogalactopyranoside (IPTG). Cultures were grown at 20°C overnight and then harvested.

All MST2 variants were purified as follows. Cells were lysed in 50mM Tris, pH 8.0, 400mM NaCl, 5% glycerol supplemented with Protease Inhibitor Cocktail (Sigma, MO). Clarified lysate was incubated with Nickel charged Profinity-IMAC resin (Biorad, CA) for one hour at 4°C, and protein eluted with 125mM imidazole. The hexa-histidine and SUMO tags were removed following incubation with SENP protease (made in house). Cleaved proteins were further purified by anion exchange and gel-filtration chromatographies. Final proteins were flash frozen at 10mg/mL in 10mM Tris, pH 8.0, 200mM NaCl, 5% glycerol, and 1mM Tris(2-carboxyethyl)phosphine (TCEP).

To generate phosphorylated MST2-K, following anion-exchange chromatography variants were incubated with phosphorylated H6-MST2-FL (made in house(8)) and 5mM ATP, 10mM MgCl_2_, 5mM NaF, 1mM Na_3_VO_4_ for 30 minutes at room temperature. Reactions were further purified on Ni-IMAC resin to remove H6-MST2-FL, and to the flow-through 20mM of ethylenediaminetetraacetic acid (EDTA) was added to quench the reaction. Phosphorylation of the variants were confirmed by Western-blot, and phosphorylated MST2-K proteins were further purified by size-exclusion chromatography.

The DNA encoding a truncated human MOB1A (residues 16-216, UniProtKB Q9H8S9) (MOB1A^τι^_15_) were cloned into a modified pBAD4 plasmid(39) downstream from N-terminal hexahistidine and SUMO tags. MOB1A^τι^_15_ was co-expressed with MBP tagged λ Phosphatase in pRSF-Duet. Plasmids were transformed into T7 express cells (New England Biolabs). Cells were grown to OD_600_ of 1.0 at 37°C, induced with 0.5mM IPTG, and grown an additional 16 hours at 20°C. Unphosphorylated MOB1A^τι^_15_ was purified following the same protocol as for unphosphorylated MST2-K.

#### CD Spectroscopy

Far-UV CD spectra were collected on an Aviv Model 410 spectrophotometer (Aviv Biomedical, NJ). CD samples contained 11μM of each MST2-K variant in 10mM Tris, pH 8.0, 200mM NaCl, 5%glycerol, and 0.5mM TCEP (ThermoFisher). Spectra were recorded at 20°C using a 0.1 cm pathlength cuvette. Data were collected with a 1nm step size, averaging for 3 seconds at each step. Before analysis, spectra from the buffer control were subtracted from each spectra; this blank subtracted data was converted to molar residue ellipticity (MRE). Data was plotted in Prism.

#### Analytical Ultracentrifugation assays

All proteins were dialyzed overnight into 10mM Tris, pH 8.0, 200mM NaCl, 5% glycerol, and 1mM TCEP. The resulting dialysate was used as a reference. Proteins were diluted in dialysate to 8.5μM, 17.1μM, 25.7μM, 34.3μM. AUC cells were assembled using SedVel60K 1.2-cm meniscus-matching centerpieces (SpinAnalytical) and sapphire windows. Sample and dialysate were loaded into the cells and spun at 25,000rpm in a Beckman XL-I until the menisci of sample and buffer matched. Concentrations were slightly diluted as a result of meniscus matching. The cells were then removed and inverted multiple times to remix the samples. The samples were allowed to return to equilibrium under vacuum at 20°C for 2hr. Sedimentation velocity experiments were run at 50,000rpm using interference optics collecting scans every 30sec for each cell for a total of 980 scans.

Sedimentation coefficient g(s*) distributions were generated from a subset of primary data where the sedimenting boundary curves had completely cleared the meniscus and located approximately in the middle of the centrifuge cell using DCCT+ by John Philo (version 2.5.1)(40,41). Partial specific volumes were calculated from the protein sequence and buffer density from the individual components using SEDNTERP(42). Data was plotted in Prism.

#### In vitro autophosphorylation assays

10μM of unphosphorylated MST2-K was incubated in 10mM Tris, pH 8.0, 150mM NaCl, 5mM MgCl_2_, 0.5mM MnCl_2_, 1mM NaF, 1mM Na_3_VO_4_, 4% glycerol, and 1mM TCEP in the presence or absence of 1mM ATP at room temperature for 60 minutes. Reactions were quenched with 20mM EDTA, and samples analyzed by Coomassie stained SDS-PAGE and Western blot using a primary antibody that recognizes MST2 phosphorylated at T180 (Cell Signaling, Lot 1, 1:1000 dilution) and the secondary antibody, IRDye 800CW Goat anti-Rabbit antibody (LI-COR, Lot C90220-06, 1:10,000 dilution). Reactions were performed three times. Band intensities were quantified, and the relative fraction of phosphorylated MST2 determined by dividing the intensity of the pT180 band by the intensity of the Coomassie band and then normalizing the signal between blots to a loading control included on each blot.

#### Cell-based autophosphorylation assays

HEK293T cells were seeded at 0.2*10^6^cells/well in 6-well plates, transfected with indicated plasmids twenty-four hours later, and lysed twenty-four hours following in ice-cold RIPA buffer supplemented with protease and phosphatase inhibitors (25mM Tris pH 7.5, 150mM NaCl, 0.2% Sodium Dodecyl Sulfate, 1% Nonidet P-40, and 1% Sodium Deoxycholate) supplemented with 1mM PMSF, 1mM Na_3_VO_4_, 10mM NaF, 2.5mM Na_2_P_2_O_7_, 1mM β-glycerophosphate, 5mM EDTA. and Protease Inhibitor Cocktail. Clarified lysates were normalized by BCA Assay and normalized samples run on Western Blot. Blots were probed with primary antibodies diluted 1:1000 that recognized either HA (Roche, Lot34502100), MYC (Santa Cruz, Lot L1318), or pT180 (Cell Signaling, Lot 2) epitopes followed secondary antibodies diluted 1:10,000 by either IRDye800CW Goat anti-Rat (LI-COR, Lot D00225-01), IRDye800CW Goat anti-Mouse(LI-COR, Lot C81106-01), or IRDye800CW goat anti-Rabbit (LI-COR, Lot C90220-06), respectively. The fraction of pT180 was determined by dividing the intensity of pT180 band by the intensity of the epitope tag and then dividing the result by the fraction of pT180 for wild-type of that replicate.

#### In vitro substrate phosphorylation assays

0.1μM of phosphorylated MST2-K was incubated with MOB1A_15_ in the same buffer used for autophosphorylation reactions in the presence or absence of 1mM ATP at room temperature for 30 minutes. Reactions were quenched with 20mM EDTA, and samples analyzed by Coomassie stained SDS-PAGE and Western blot using a primary antibody that recognizes phosphorylated T35 of MOB1A (Cell Signaling, Lot 2, 1:1000 dilution) followed by IRDye 680RD goat anti-rabbit antibody (LI-COR, Lot C71115-11, 1:10,000 dilution). Reactions were performed three times. Band intensities were quantified, and the relative fraction of phosphorylated MOB1A_15_ determined by dividing the intensity of the pT35 band by the intensity of the Coomassie band and normalizing to the fraction pT35 for wild-type MST2-K.

#### General Tissue Culture

HEK293T cells (ATCC, VA) were cultured in DMEM:F12 medium (Gibco) supplemented with 5% Fetal Bovine Serum (VWR) and 2mM Glutamine (Gibco) and grown at 37°C in 5%CO_2_. Cells were transfected with indicated plasmids using polyethyleimine “MAX” (Polysciences)(43).

#### Image Quantification and Statistical Analysis

Both Coomassie stained gels and Western blots were scanned on an Odyssey Infrared Imaging System (LI-COR) barring avidity immunoprecipitations of MST2 variants in Figures 4,5 and Figure S2 which were imaged on a GBox (Syngene). Band intensities were quantified using ImageJ(38). Statistical analysis and plotting of data was performed in Prism (GraphPad software, La Jolla California, USA).

## DATA AVAILABILITY

Requests for raw data, additional information, or reagents contained within the manuscript are available upon request from Jennifer Kavran, Johns Hopkins Bloomberg School of Public Health.

## SUPPORTING INFORMATION

This article contains supporting information.

## ACKNOWELDGEMENTS

We thank the Johns Hopkins University Center for Molecular Biophysics for providing instrumentation and resources and David Snead for critical reading of the manuscript.

## FUNDING AND ADDITIONAL INFORMATION

This work is supported by NIH R01GM134000 to JMK and NIH T32CA009110 for TT and KW. The content is solely the responsibility of the authors and does not necessarily represent the official views of the National Institutes of Health.

## ABBREVIATIONS

(COMP): cartilage oligomeric matrix protein
(CD): Circular dichroism
(IPTG): Isopropyl β-d-1-thiogalactopyranoside
(MBP): Maltose binding protein
(PMSF): Phenylmethylsulfonyl fluoride
(TCEP): 1mM Tris(2-carboxyethyl)phosphine
(EDTA): ethylenediaminetetraacetic acid
(AMP-PNP): Adenylyl-imidodiphosphate
(HRP): Horseradish peroxidase
(AP): Alkaline phosphatase
(MRE): Molar residue ellipticity
(RMSD): Root-mean-square deviation
(K_d_): Dissociation constant
(MST): Mammalian sterile twenty-like
(EGFR): Epidermal growth factor receptor
(GCK): Germinal center kinases
(AUC): Analytical Ultracentrifugation

## Supporting Information

### Table of Contents

**Figure S1**. *Structure gallery of MST kinase dimers*.

**Figure S2**. *Proof of concept for avidity pull downs*

**Figure S3**. *Replicates for avidity pull downs*

**Table S1**. *Pairwise RMSD for chain A onto chain A for all kinase-domain pairs*

**Table S2**. *Pairwise RMSD for chain A onto chain B for all kinase-domain pairs*

**Table S3**. *Analysis of all kinase-domain pairs*

**Figure S1.**
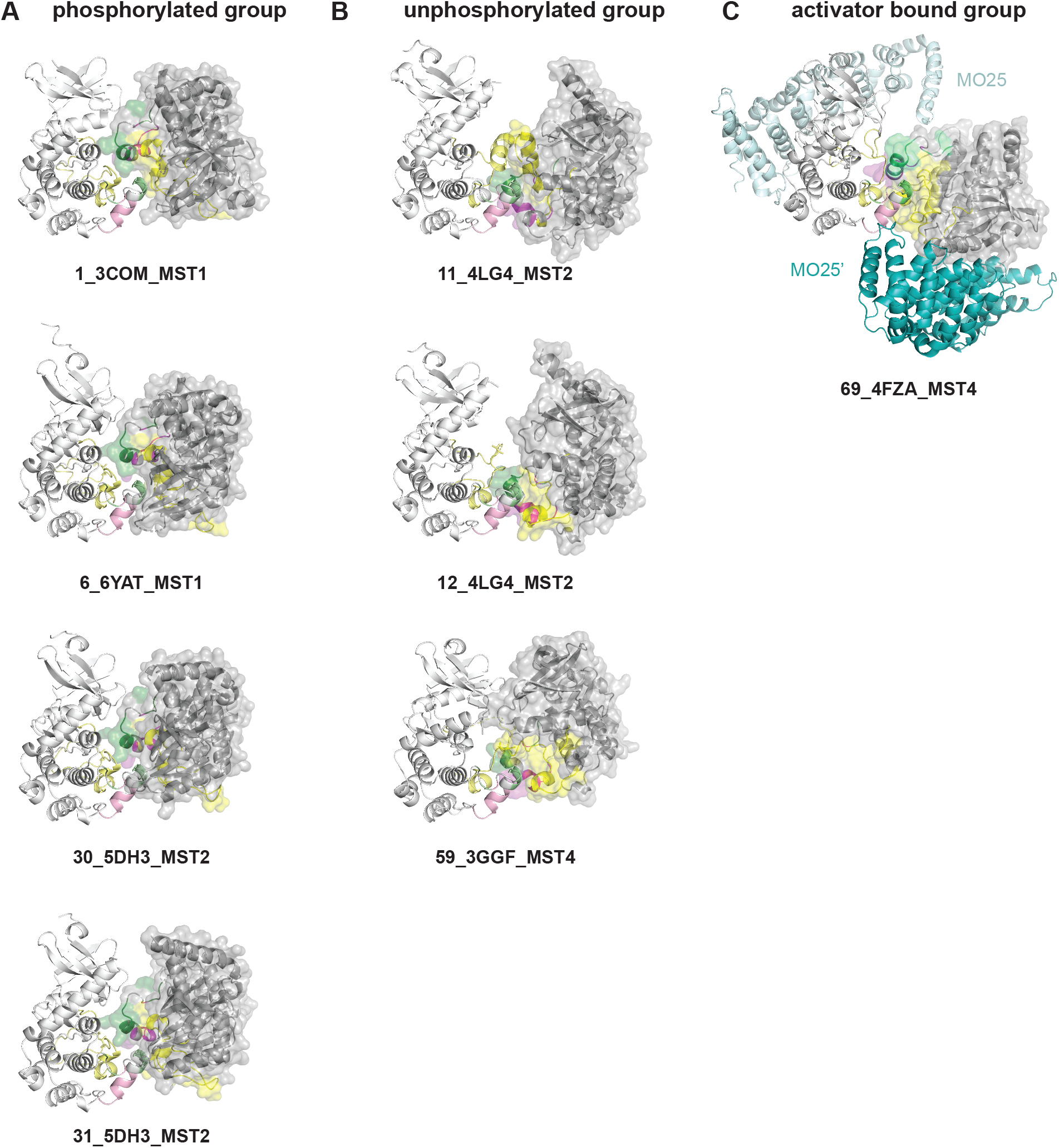
Gallery of MST2 kinase domain dimers compatible with trans-autophosphorylation. For each dimer one chain is shown in white cartoon and the other in dark gray cartoon and a semi-transparent surface. The residues that comprise GC and GN are shown in pink and green, respectively. The activation loop is indicated in yellow. All unphosphorylated dimers are in (A), phosphorylated dimers in (B), and bound to the activator MO25 (teal) (C).

**Figure S2.**
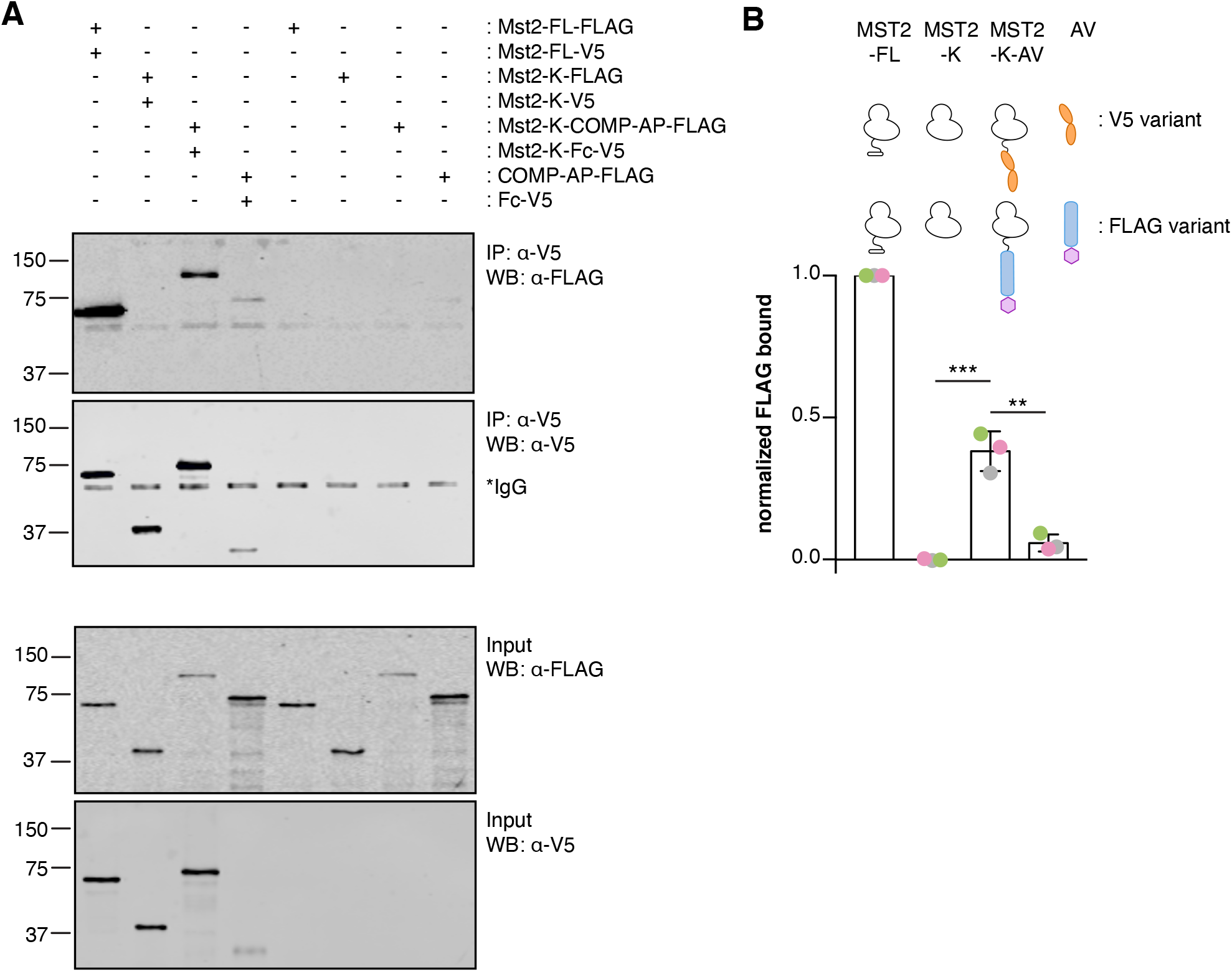
Avidity-based immunoprecipitation of MST2 kinase domains. HEK293 cells transiently transfected as indicated, and complexes isolated from lysates following immunoprecipitation with V5 antibody. Protein expression and complex formation were monitored by Western blot. (A) A representative set of blots are shown. An asterisk indicates a band corresponding to IgG used in the pull-down. (**B**) Band intensities of bound FLAG-tagged proteins were quantified ((bound-background)/normalization). The bar graph represents the mean from three experiments, and the errors bars the standard deviation. A cartoon below each bar represents the proteins used. Significant differences were calculated using an unpaired *t*-test (**, *p* ≤ 0.002; ***, *p* ≤ 0.0008).

**Figure S3.**
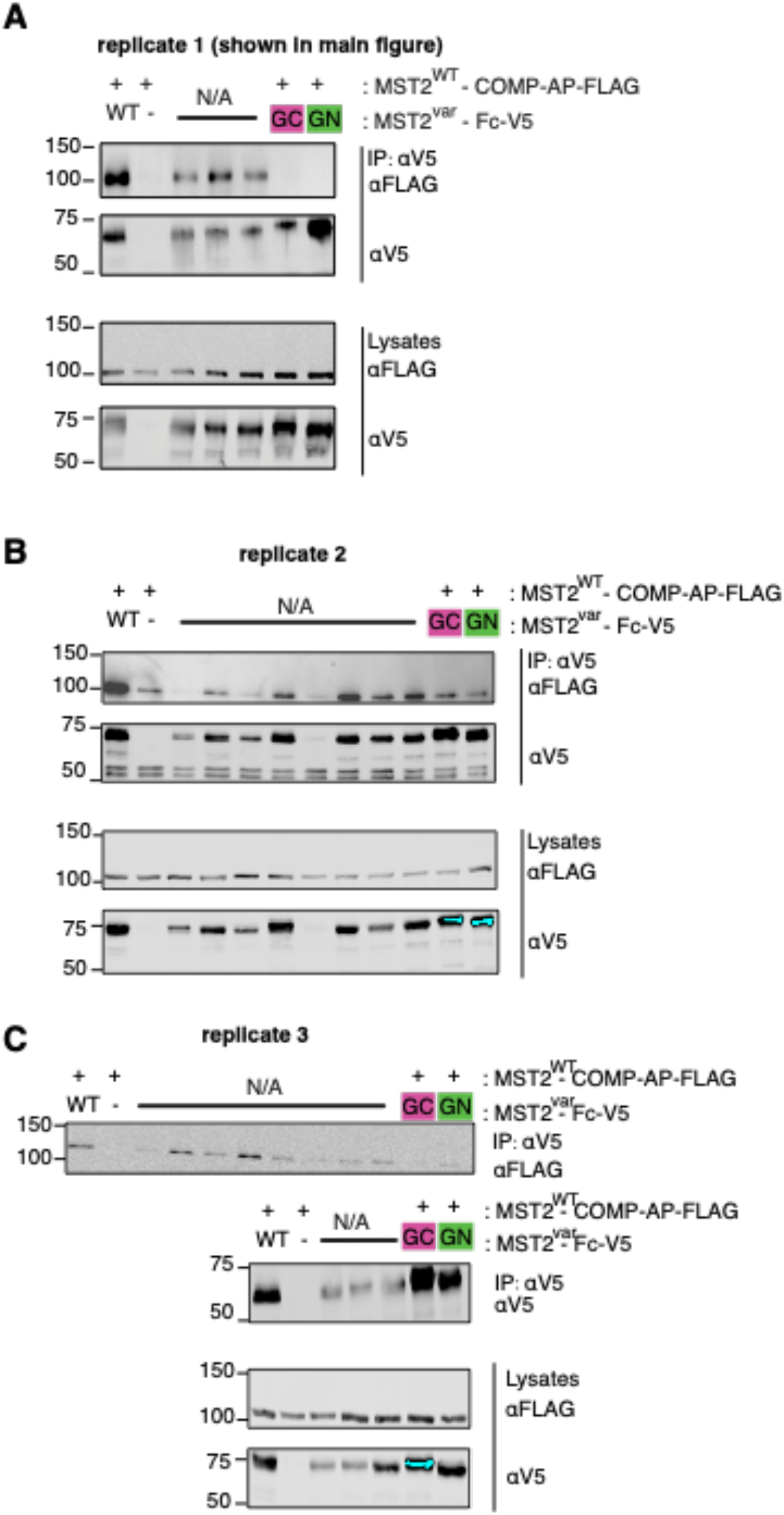
Replicates of avidity pull downs. Three replicates of avidity-based immunoprecipitations, including uncropped blots, used for quantification of Figure 4. Lanes not relevant to the presented study are indicated by “N/A”, not applicable.

**Table S1.**
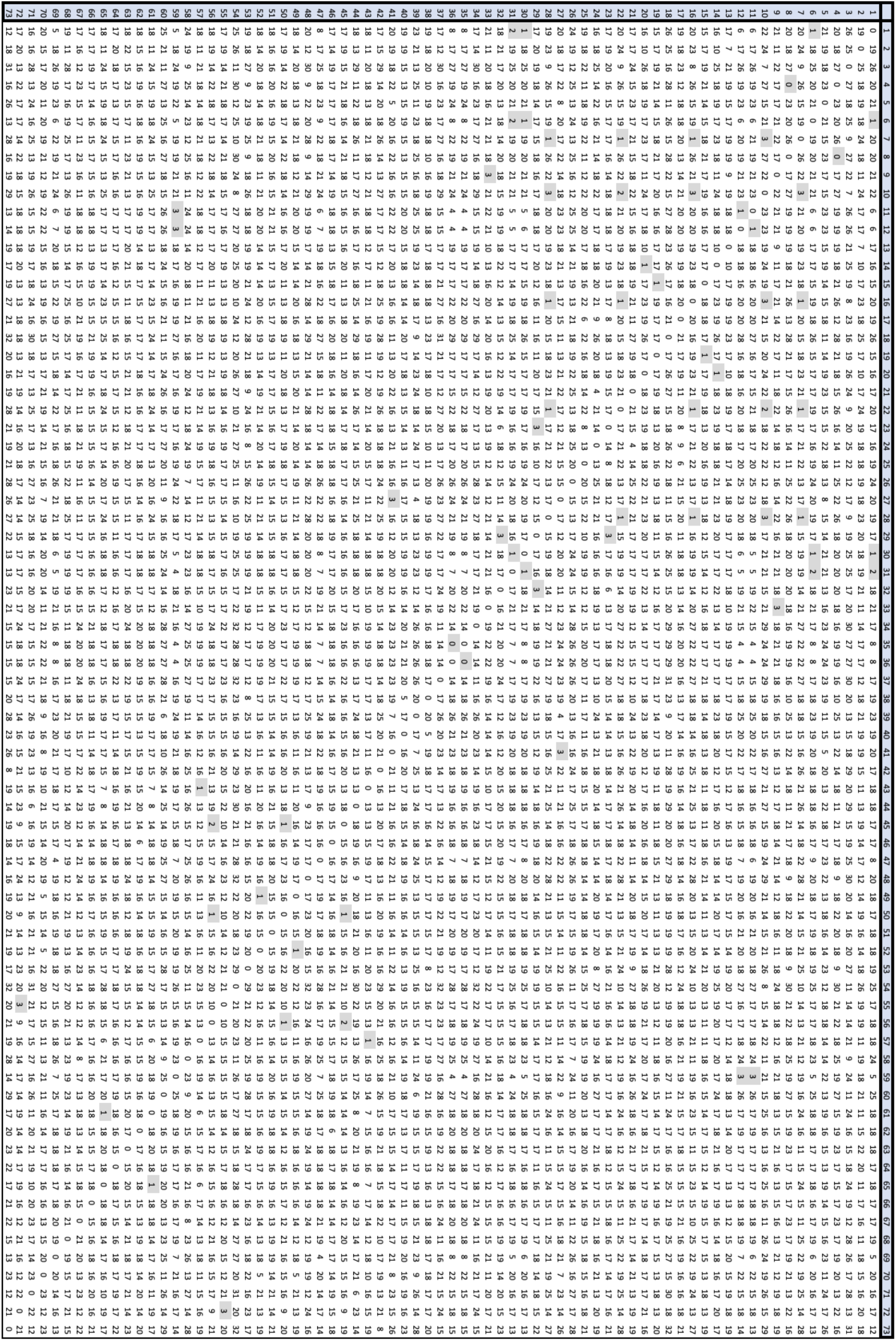
Pairwise RMSD for A onto A for all potential kinase-domain dimers ^a^ values reported in Å ^*b*^ kinase-domain naming abbreviated to number only, these names are highlighted in blue ^c^ RMSD less than 3Å are highlighted in gray

**Table S2.**
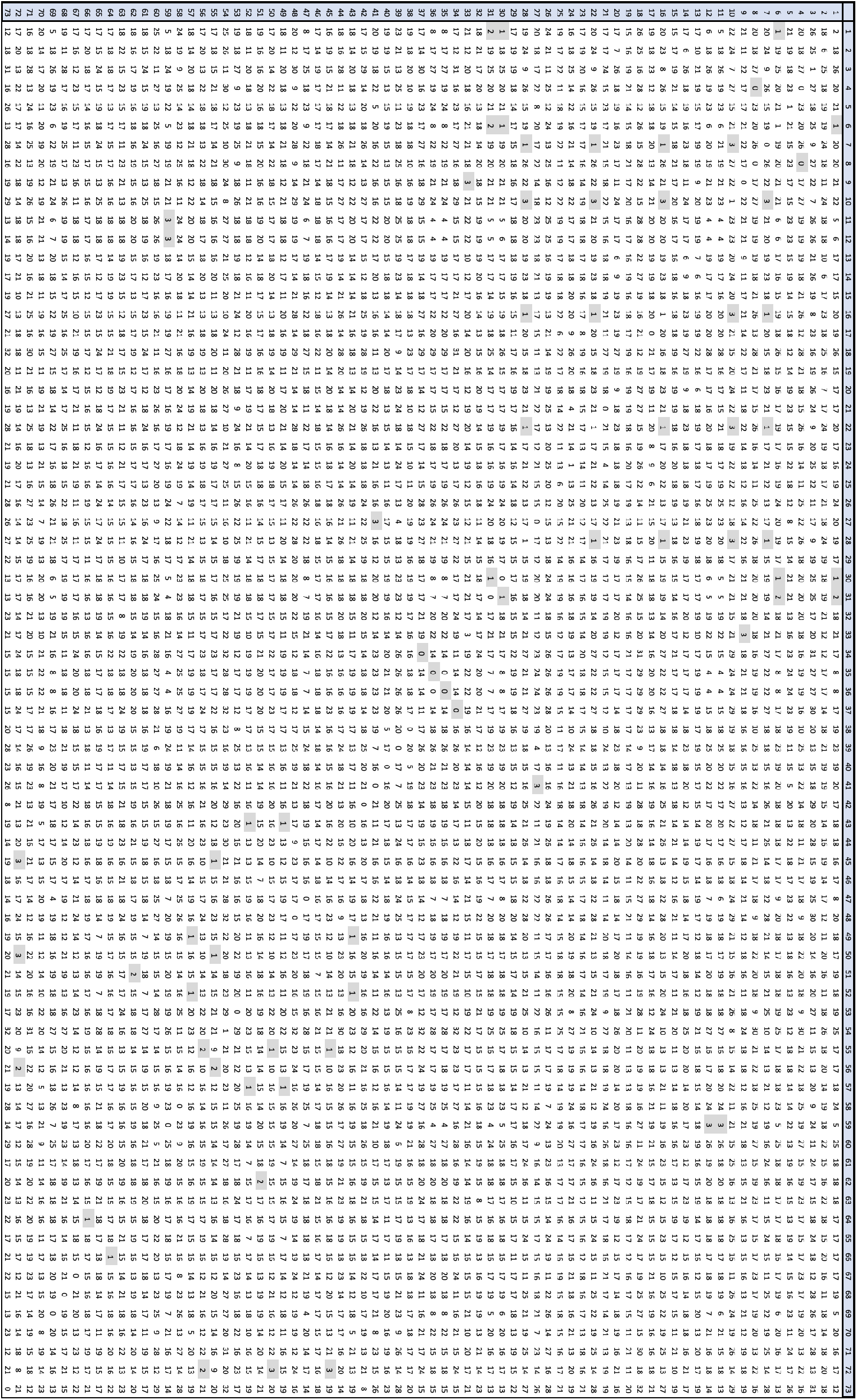
Pairwise RMSD for A onto B for all potential kinase-domain dimers ^a^values reported in Å ^b^ kinase-domain naming abbreviated to number only, these names are highlighted in blue ^c^ RMSD less than 3Å are highlighted in gray

**Table S3.**
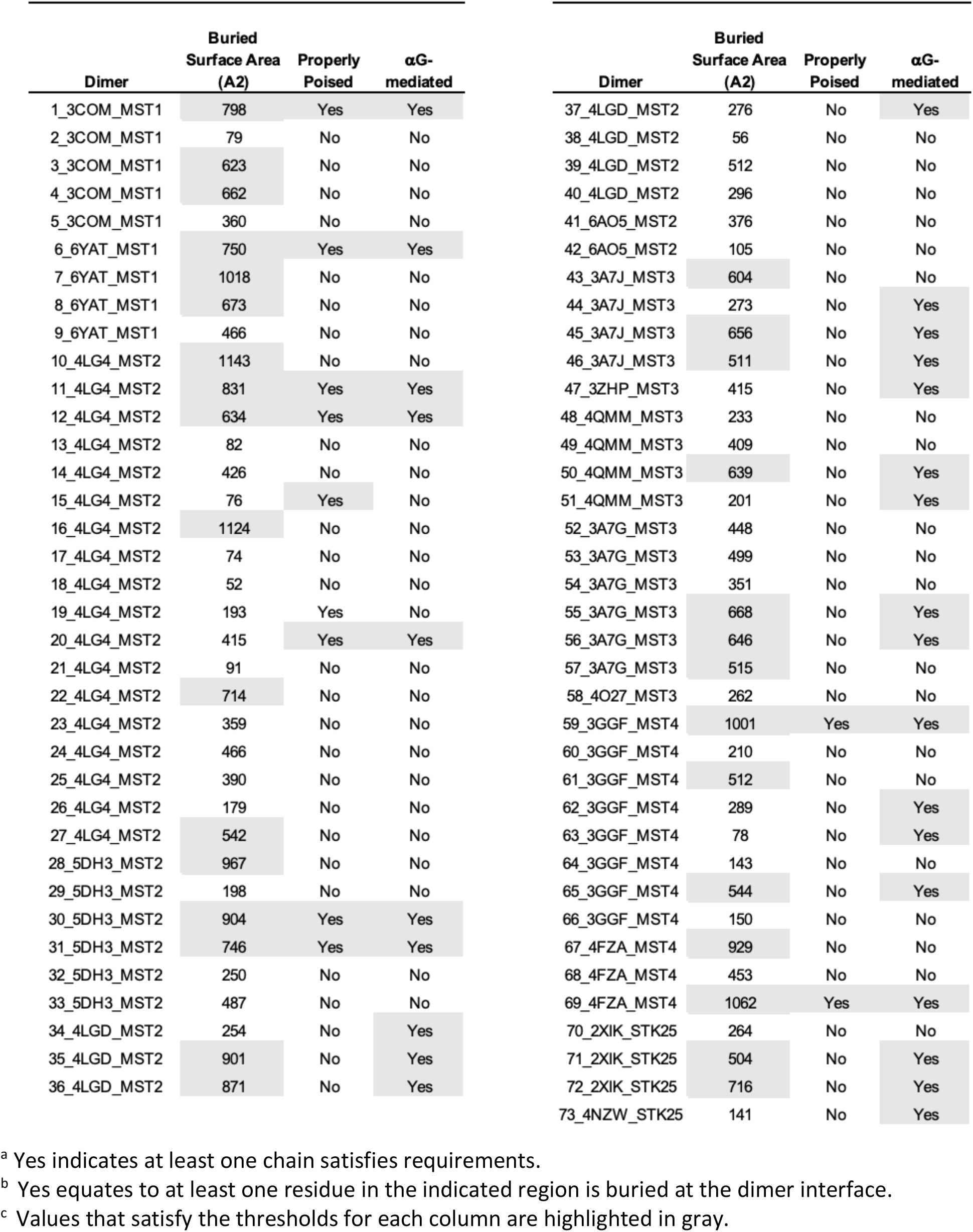
Analysis of all kinase-domain pairs

